# Growth temperature is the principal driver of chromatinization in archaea

**DOI:** 10.1101/2021.07.08.451601

**Authors:** Antoine Hocher, Guillaume Borrel, Khaled Fadhlaoui, Jean-François Brugère, Simonetta Gribaldo, Tobias Warnecke

## Abstract

Across the tree of life, DNA in living cells is associated with proteins that coat chromosomes, constrain their structure and influence DNA-templated processes such as transcription and replication. In bacteria and eukaryotes, HU and histones, respectively, are the principal constituents of chromatin, with few exceptions. Archaea, in contrast, have more diverse repertoires of nucleoid-associated proteins (NAPs). The evolutionary and ecological drivers behind this diversity are poorly understood. Here, we combine a systematic phylogenomic survey of known and predicted NAPs with quantitative protein abundance data to shed light on the forces governing the evolution of archaeal chromatin. Our survey highlights the Diaforarchaea as a hotbed of NAP innovation and turnover. Loss of histones and Alba in the ancestor of this clade was followed by multiple lineage-specific horizontal acquisitions of DNA-binding proteins from other prokaryotes. Intriguingly, we find that one family of Diaforarchaea, the Methanomethylophilaceae, lacks any known NAPs. Comparative analysis of quantitative proteomics data across a panel of 19 archaea revealed that investment in NAP production varies over two orders of magnitude, from <0.02% to >5% of total protein. Integrating genomic and ecological data, we demonstrate that growth temperature is an excellent predictor of relative NAP investment across archaea. Our results suggest that high levels of chromatinization have evolved as a mechanism to prevent uncontrolled helix opening and runaway denaturation – rather than, for example, to globally orchestrate gene expression – with implications for the origin of chromatin in both archaea and eukaryotes.

## INTRODUCTION

Archaeal genomes are coated by small, abundant and often basic proteins that bind DNA with low sequence specificity. As in bacteria, these are collectively referred to as nucleoid-associated proteins (NAPs). A variety of proteins that fit this loose description have been described in archaeal model organisms, and include histones, Alba, Cren7, and MC1 (Zhang *et al*. 2012). Whereas histones provide the backbone of chromatin across eukaryotes, the repertoire of major chromatin proteins in archaea is considerably more diverse and fluid. Histones are common but were lost along several lineages, including the Sulfolobales/Desulfurococcales and Parvarchaeota (Adam *et al*. 2017). On the flipside, several NAPs are abundant but lineage-specific, including HTa in the Thermoplasmatales (Hocher *et al*. 2019) and Sul7 in the Sulfolobales (Zhang *et al*. 2012).

The evolutionary and ecological drivers of NAP turnover in archaea are poorly understood. In fact, the diversity of NAPs in archaea has been described as “puzzling” in light of otherwise highly conserved information processing pathways (Visone *et al*. 2014). Do different NAPs represent adaptations to specific niches? If so, what factors determine the presence or absence of a given NAP in a given genome? Alternatively, are NAPs in archaea diverse because several different proteins can do the same job, rendering them exchangeable? In other words, are these NAPs functional analogues that can replace each other without serious repercussions for genome function?

Phylogenomic surveys provide a useful starting point to tackle these questions. They chart the distribution of homologous genes across a set of genomes and allow gain and loss events to be traced along the phylogeny. The resulting presence/absence patterns, considered in ecological context, might then reveal clues as to why a particular gene is found in one set of genomes but not another. Phyletic comparisons, however, can be treacherous. The presence of a specific gene in two genomes does not necessarily imply that the protein product is doing the same job in both. Indeed, in the case of NAPs, we have reason to suspect that this assumption – known as the ortholog conjecture – is not always met. Histones, for example, are highly abundant at the protein level in *Thermococcus kodakarensis* (1.76% of total protein) but only weakly expressed in *Halobacterium salinarum* (0.02% of total protein) (Rojec *et al*. 2019; Müller *et al*. 2020). Given the difference in abundance, histones are unlikely to play the same roles in nucleoid biology in these two species. In line with this, the single *H. salinarum* histone gene (*hpyA*) is dispensable for growth (Dulmage *et al*. 2015), whereas retention of at least one of its two histone genes (*htkA, htkB*) is essential in *T. kodakarensis* (Cubonova *et al*. 2012). Alba too is highly expressed in many archaea, including *Sulfolobus shibatae* (1.6% of total protein) but >100-fold less abundant in others, e.g. *Methanococcus maripaludis* (0.01% of total protein) (Liu *et al*. 2009). These large differences in abundance are indicative of cryptic functional diversity that is not directly accessible via comparative genomics.

Here, we combine a systematic phylogenomic survey of NAPs with quantitative mass spectrometry data on NAP abundance to shed light on the evolutionary drivers of chromatin diversity in archaea. Our survey highlights the Diaforarchaea as a group that has experienced exceptional NAP fluidity. Most surprisingly, we find that the Methanomethylophilaceae, a family in the order Methanomassiliicoccales, are devoid of any known NAPs. We generate quantitative proteomics data for two members of the Methanomassiliicoccales, *Methanomethylophilus alvus* and *Methanomassiliicoccus luminyensis*, and develop a simple pipeline to identify novel NAP candidates in these species. Applying this pipeline to proteomic data from an additional 17 archaea, we propose several new proteins with the capacity to play global roles in nucleoid organization, including highly abundant but previously uncharacterized proteins in model archaea. Pan-archaeal analysis revealed substantial quantitative variation in NAP abundance. Total investment in NAPs varies over two orders of magnitude, from as little as 0.014% to 5.38% of total protein. Exploring potential ecological and genomic covariates of differential investment in NAPs, we pinpoint growth temperature as the principal driver of NAP abundance across archaea. Our results suggest that high levels of chromatinization in archaea are first and foremost an adaptation to thermal stress. We speculate that chromatin in eukaryotes might constitute an evolutionary hangover from their thermophilic past: unlike many of their archaeal cousins, eukaryotes were unable to reduce their large investment in chromatin proteins as they transitioned to a mesophilic lifestyle, presumably because histones had assumed important functions other than thermoprotection.

## RESULTS

### A phylogenomic survey uncovers archaea without known nucleoid-associated proteins

To provide an up-to-date view of NAP diversity across the archaea, we first collated a list of previously described archaeal NAPs (Fig 1A) and used sensitive Hidden Markov Model (HMM) scans to establish the presence/absence of NAP homologs in 1419 archaeal genomes representative of known archaeal diversity (Table S1, see Methods). As highlighted previously (Zhang *et al*. 2012), archaeal chromatin is not dominated by a single protein but by small cliques of typically two (and sometimes three or more) abundant proteins (Fig 1B/C, Table S1). Different NAPs from a pan-archaeal repertoire can co-occur in most any clique, which are frequently dismantled by gene loss and absorb new members via horizontal gene transfer (HGT). Across our sample, any given NAP can be found partnering any other (Fig 1B, Fig S1), suggestive of functional promiscuity. While some NAPs are phylogenetically widespread, none are universal to archaea (Fig 1B/C). Histones and Alba are the most common and were likely present in the last archaeal common ancestor (LACA), but both have been lost along different lineages (Fig 1C). Conversely, gains are common and frequently driven by HGT (see below).

**Figure 1.**
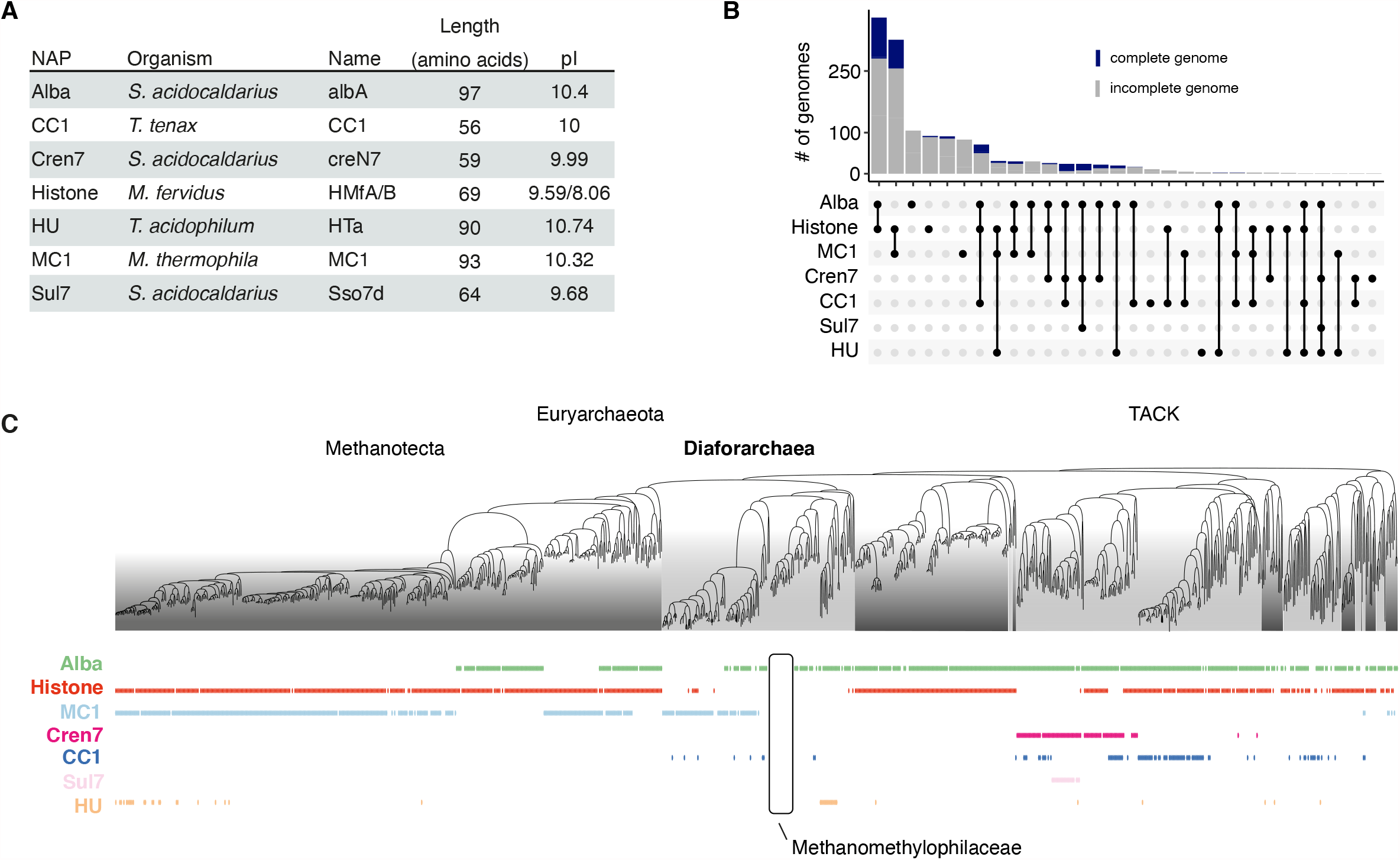
Distribution of nucleoid-associated proteins (NAPs) across archaea. (**A**) Names and properties of previously characterized archaeal NAPs and (**B**) their co-occurrence in 1419 sequenced archaeal genomes. (**C**) Presence/absence of NAPs in phylogenetic context, highlighting the Methanomethylophilaceae as a family without any previously characterized NAPs. The species-level phylogeny is based on GTDB (see Methods). See Table S1 and Fig S1 for species-level information.

One clade where fluidity in NAP repertoire is particularly striking is the Diaforarchaea (Fig 2A). Both histones and Alba were lost at the root of this clade. Several lineages, including the Thermoplasmatales, later re-acquired Alba from different sources, as supported by the polyphyletic distribution of diaforarchaeal homologs on a pan-archaeal Alba tree (Fig 2B). Subsequent to *alba*/histone loss, NAP repertoires evolved in a highly idiosyncratic fashion along different diafoarchaeal lineages. We previously described the presence of an HU homolog (HTa) in *Thermoplasma acidophilum*, which is highly expressed, exhibits histone-like binding behaviour, and was likely acquired via HGT from bacteria (Searcy 1975; Hocher *et al*. 2019). HU homologs, however, are confined to the Thermoplasmatales and absent from the remainder of the Diaforarchaea (Fig 2A). Similarly, most members of the marine group II (MG-II) archaea encode MC1, a NAP best known from Methanosarcina *spp*. (Chartier *et al*. 1988) and widespread amongst haloarchaea (Fig 1C). Again, MC1 is only present in MG-II but absent from other diaforarchaeal lineages, and was likely acquired via HGT (Fig 2A, S2). Most curiously, we find that the members of one diaforarchaeal lineage, the Methanomethylophilaceae, encode no known NAPs whatsoever (Fig 2A).

**Figure 2.**
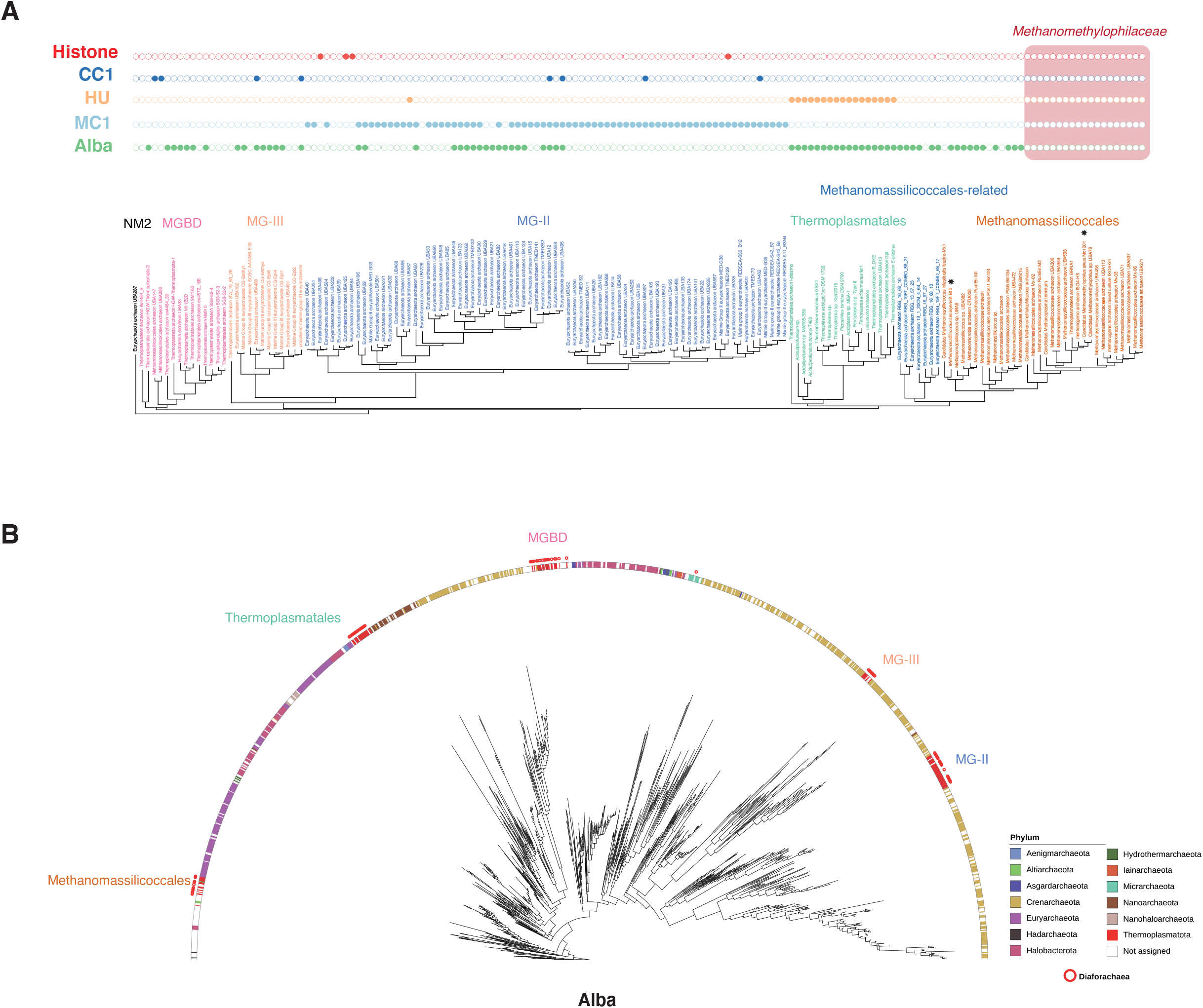
Nucleoid-associated proteins (NAPs) in the Diaforarchaea. (**A**) Presence/absence of NAPs in phylogenetic context, highlighting the absence of known NAPs in the Methanomethylophilaceae, the lineage-restricted distribution of HU and MC1, and the patchy distribution of Alba. The species-level phylogeny is based on GTDB (see Methods). The two species for which proteomics data were generated are marked with asterisks. (**B**) Maximum likelihood protein tree (see Methods) of archaeal Alba homologs. Alba sequences from the Diaforarchaea are not monophyletic, suggesting multiple independent acquisition events. MG-II (III): Marine group II (III); NM2: New Methanogen lineage 2; MGBD: Marine benthic group D.

### Predicting novel candidate NAPs

Do members of the Methanomethylophilaceae make do without major nucleoid-associated proteins? Or do these organisms encode as yet uncharacterized NAPs? To begin to address these questions, we generated quantitative mass spectrometry data for two members of the Methanomassiliicoccales, both isolated from the human gut: *Methanomassiliicoccus luminyensis* (Dridi *et al*. 2012) and *Methanomethylophilus alvus* (Fadhlaoui, Ben Hania *et al*., in preparation). We detected and quantified 72% of the predicted proteome in *M. alvus* and 67% in *M. luminyensis*, which compares favourably to a recent state-of-the-art effort to catalogue proteins across the tree of life (Müller *et al*. 2020) (Fig S3). Alba, though present in the genome of *M. luminyensis*, was not expressed at detectable levels. We then developed a simple pipeline to predict proteins that might play a global role in nucleoid organization similar to known NAPs (Fig 3A, Methods). To qualify as a candidate NAP, proteins had to meet four criteria: First, they could not exceed the size of previously characterized NAPs, which are typically short. Permissively, we only considered proteins smaller than 290 amino acids, 110% the size of TrmBL2 in *T. kodakarensis* (see below). Second, they had to contain either a known DNA-binding domain or be predicted to bind DNA (see Methods). Third, they needed to be expressed at a level that makes them high-abundance outliers compared to other predicted transcription factors, objectively determined using Rosner tests (see Methods). Fourth, they had to be encoded as single-gene operons. The latter criterion was adopted following the observation that known NAPs are usually present in single-gene operons (Fig 3B, see Methods). Two proteins in *M. luminyensis* met these criteria. One of them (WP_019177984.1) is a small (74 amino acids), basic (pI: 9.64) and lysine-rich protein, which constitutes 1.34% of the *M. luminyensis* proteome (Fig 3C, Table S1), making it the 12^th^ most highly expressed protein in our sample. Its homolog in *M. alvus* (AGI86273.1) was independently identified as the sole candidate NAP in this organism, where it is less strongly expressed (0.14% of total protein, ranking 123^rd^ out of 1220 proteins). Neither protein contains a known DNA-binding domain. Phylogenomic analysis reveals orthologs of WP_019177984.1/AGI86273.1 throughout the Methanomassiliicoccales and in several bacterial genomes, particularly amongst Sphingomonadales and Rhodobacterales (Fig S4). Monophyly of the Methanomassiliicoccales homologs suggests a single acquisition event at the root of this clade, which preceded the loss of Alba (Fig S4, Fig 2). Experimental investigation will be required to confirm or refute the association of WP_019177984.1/AGI86273.1 with the nucleoid, but we suggest – based on this preliminary evidence – that it is unlikely that Methanomethylophilaceae make do entirely without NAPs. Relaxing criteria on single-gene operon status did not reveal additional candidates for *M. alvus*. Two additional candidates were recovered in *M. luminyensis* but their quantitative contribution to overall NAP investment is minor (0.4%).

**Figure 3.**
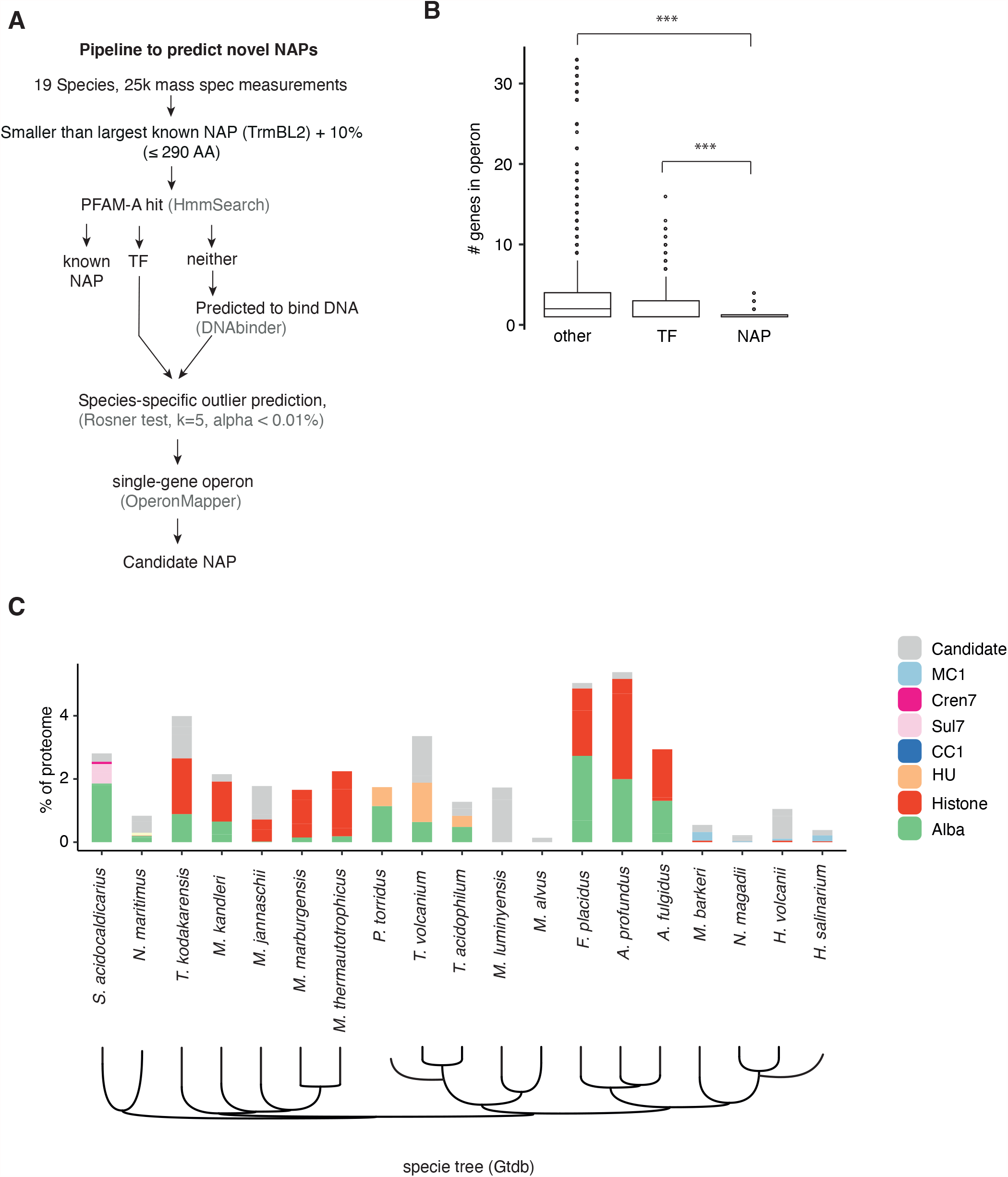
Quantitative variation in the abundance of known and predicted nucleoid-associated proteins (NAPs). (**A**) Outline of the bioinformatic pipeline to predict novel NAPs. Proteins detected by mass spectrometry need to pass several successive filters to be considered as a candidate NAP (see main text and Methods for a detailed description of the individual steps). (**B**) Sizes (number of genes) for operons that contain proteins classified as NAPs, transcription factors (TF) or “other” domains (neither NAP nor TF) based on Pfam annotations. (**C**) Large variation in the relative abundance (% of proteome) of known and candidate NAPs in 19 species of archaea for which quantitative mass spectrometry data was analyzed. The species tree is taken from GTDB, with *P. torridus* and *H. salinarum* added manually.

### Detection of novel candidate NAPs in several model archaea

To see whether the same approach might reveal additional candidate NAPs in other archaea, including well-characterized model species, we applied the same pipeline to a compendium of 17 archaea for which we could retrieve published proteome-scale quantitative mass spectrometry data. Quantitative inventories for 13 of these species were recently published as part of a cross-kingdom proteome survey (Müller *et al*. 2020) and generated using the same protocol that we used for *M. luminyensis* and *M. alvus* (see Methods).

Starting from 22643 proteins across 17 species, and excluding known NAPs, we retrieved 22 candidate hits (Fig 3D; Fig S5). Manual inspection revealed one obvious false positive, AAY80109.1 in *Sulfolobus acidocaldarius*, a truncated Lrp/AsnC transcriptional regulator that, unlike in other Sulfolobales, lacks a DNA-binding domain. Reassuringly, we recover TrmBL2, a known constituent of chromatin in *T. kodakarensis* where it is unusually abundant compared to TrmB homologs in other archaea (Fig S6). For some species (e.g., *Methanothermobacter marburgensis* and *Archaeoglobus fulgidus*) we identified no additional candidates, suggesting that our pipeline is not excessively greedy (Fig 3D). For others (e.g., *Sulfolobus acidocaldarius*), we only retrieved candidates that are much less abundant than known NAPs in the same organism. In contrast, we also observed species where novel candidates make up a substantial portion of the overall investment in NAPs, rivalling or even dwarfing the abundance of known NAPs. Notably, this list includes the model archaeon *Haloferax volcanii*, where the two candidate NAPs (HVO_1577, HVO_2029) are considerably more abundant than either histones or MC1 (Fig 3D), a finding we confirm in an independently generated proteomics dataset (Fig S7). We note that, unlike the NAP candidate WP_019177984.1/AGI86273.1 in *M. luminyensis/M. alvus*, most of these hits are larger than well-known NAPs and might therefore, like TrmBL2, have originated from normally less abundant transcription factors.

### Investment in NAPs varies extensively across archaea

With or without candidate NAPs, the comparison above revealed striking variation in total NAP investment across species, ranging over two orders of magnitude, from 0.014% of total protein in the halophile *Natrialba magadii* to 5.38% in *Archaeoglobus profundus* (Fig 3D). Individual NAPs can vary over a similar range: relative histone abundance, for example, varies up to 400-fold (Fig S8), from 3.2% of the proteome in *A. profundus* to <0.06% in *Nitrosopumilus maritimus, Methanosarcina barkeri* and the Halobacteriales, where abundance is indistinguishable from that of sequence-specific transcription factors (Fig S8). Note here that we operationally define % total protein as the fraction of total intensity in the mass spectrometry data attributable to NAPs.

What are the reasons for this wide variability in NAP investment? We initially considered the possibility that variability is not, in fact, biologically meaningful but attributable to experimental factors. It is conceivable, for example, that a NAP, once detected, might represent an artificially high proportion of a proteome simply because comparatively few proteins were quantified. This is clearly not the case in *H. volcanii*: less than 60% of protein-coding genes were quantified in this species (Fig S3) but histones make up only a small proportion of this total. It is also not generally true: we find no correlation between fractional coverage of the predicted proteome and the proportion allocated to NAPs (rho=-0.31, P=0.19). Further, differential investment is also evident when relative abundance is scaled not to the total proteome but to the abundance of house-keeping genes (tRNA synthetases) that show low cross-species variability (Fig S9, see Methods).

Next, to confirm that fractional protein abundances can be reasonably compared across species, we asked whether the relative abundance of a protein in one species is usually predictive of the relative abundance of its homolog in another species. Considering reciprocal best-blast hits (RBBHs) between species as an indicator of homology, we find this to be the case. Organisms that are phylogenetically or ecologically close tend to have more correlated abundance profiles (Fig S10). This is particularly evident when we consider not individual RBBHs but instead aggregate protein abundance by Pfam domain content or gene ontology category (see Methods, Fig S10). These results indicate that we can make informative quantitative comparison across species using the proportion of total protein as a metric. The results also advocate the use of lower granularity. Below, we therefore consider the abundance of all NAPs collectively.

### Growth temperature predicts NAP investment

We next asked whether relative NAP investment is a (somewhat trivial) function of genome size, whereby organisms with larger genomes need to make a greater relative investment in NAPs because they have more DNA to coat. We find this not to be the case (rho=-0.3, P=0.21). To gain clues into potential ecological drivers of NAP investment, we then determined which proteins (or protein domains/functional categories) quantitatively covaried with NAP investment across species (see Methods). Amongst the most highly correlated domains, we find several that are classically associated with heat stress, including the protein chaperones Hsp20 and prefoldin but also RTCB, which has been implicated in recovery from stress-induced RNA damage (Tanaka and Shuman 2011) (Fig 4A). Prompted by these findings, we examined several environmental and phenotypic variables, including optimal growth temperature (OGT), pH and doubling time. We find that relative abundance of NAPs is uniquely, and strongly, associated with OGT (rho=0.83, P=8e-6, Fig 4B, Table S2). This finding is robust to inclusion/exclusion of candidate NAPs (Fig S11) and holds true for individual NAPs, notably histones and Alba, which are sufficiently widespread to allow cross-species comparisons (Fig S11). Importantly, the relationship between NAP abundance and OGT is preserved when controlling for phylogenetic non-independence (see Methods). The relative abundance of sequence-specific transcription factors, on the other hand, does not covary with temperature (Fig 4C).

**Figure 4.**
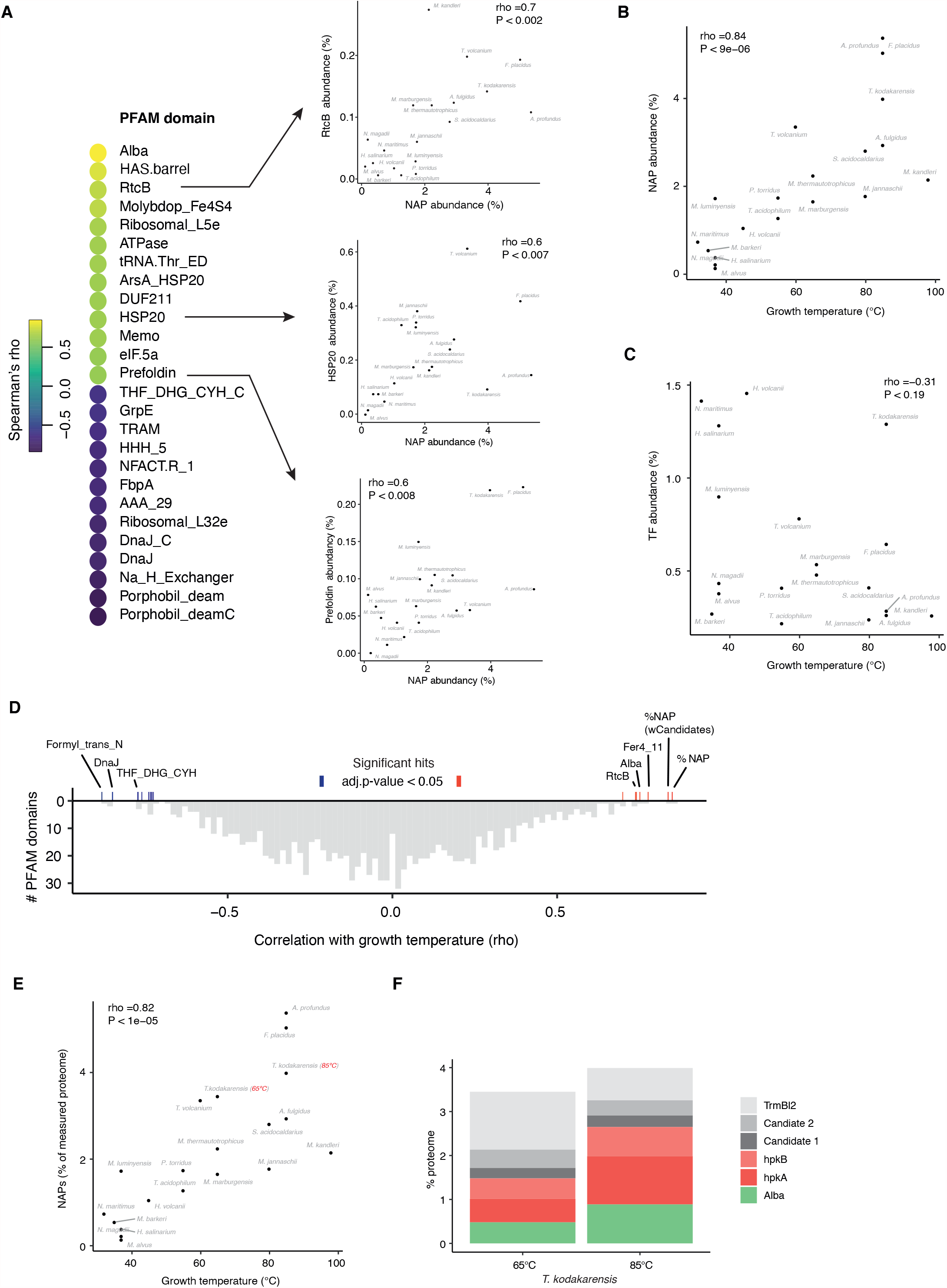
The relationship between growth temperature and NAP investment across archaea. (**A**) Top (bottom) 13 Pfam domains whose relative abundance is most positively (negatively) correlated with relative NAP abundance (aggregated across known and candidate NAPs) across the sample of 19 archaea shown in Fig 3C. Examples of individual correlations are shown on the right. (**B**) Relationship between NAP abundance and optimal growth temperature across the same set of archaea. (**C**) The aggregate abundance of transcription factors (TF) is not correlated with optimal growth temperature. (**D**) Distribution of correlation coefficients between growth temperature and the relative abundance of 1154 Pfam domains. The relative abundance of NAPs, considered as an aggregate class, exhibits the strongest correlation with growth temperature. We obtain similar results when considering gene ontology categories instead of Pfam domains. (**E, F**) Relative NAP investment drops when *T. kodakarensis* is grown at 65°C instead of its optimum growth temperature of 85°C.

Is it unexpected to find a strong relationship between the abundance of a certain class of proteins and OGT? To address this question, we computed the correlation between OGT and the relative abundance of 1154 Pfam domains (and 297 gene ontology categories) across the 19 species in our analysis. Strikingly, NAPs (considered as an aggregate class) have the strongest relationship with growth temperature (Fig 4D).

### NAP investment increases with temperature on physiological time scales

If growth temperature is a driver of NAP abundance over evolutionary time scales, this might also be true for physiological time scales. Do we see NAP abundance go up following heat stress and decrease in response to cold shock? Although no systematic data exist that span the diversity of species examined above, prior temperature shift experiments from various archaea support this hypothesis: first, NAP abundance is affected by temperature in *T. kodakarensis* (Fig 4E) (Sas-Chen *et al*. 2020), with reduced levels of histones and Alba driving a 13.5% relative drop in total NAP investment at 65°C compared 85°C (Fig 4F). Similarly, histone transcripts are downregulated upon cold shock in *M. jannaschii* (Boonyaratanakornkit *et al*. 2005), as are histones and MC1 in *Methanococcoides burtonii* (Campanaro *et al*. 2011), while Sul7 expression increases upon heat shock in *Sulfolobus solfataricus* (Tachdjian and Kelly 2006). Taken together, these observations suggest that NAP investment varies with growth temperature not only over evolutionary but also physiological time scales.

## DISCUSSION

Temperature is a powerful driving force for molecular evolution. Selection for increased thermostability has left conspicuous footprints on the composition of proteins and RNAs in many species. Proteins from thermophiles are, for example, enriched in charged and hydrophobic amino acids while their structural RNAs (tRNAs, rRNAs) exhibit higher than average GC content, consistent with the need for stronger base-pair bonds at higher temperatures (Hickey and Singer 2004; Zeldovich *et al*. 2007). Similar compositional hard-coding was suspected to occur at the DNA level: as G-C base-pairs provide greater stability than A-T pairs, it was reasonable to suspect that thermophiles have genomes with elevated GC content. However, this turned out not to be the case (Hickey and Singer 2004). As demonstrated, for example, by *T. kodakarensis* (52% GC, OGT: 85°C) and *Pyrococcus furiosus* (41% GC, OGT: 100°C), average or even low genomic GC content is clearly compatible with growth at high temperatures (Grogan 1998).

NAPs, along with polyamines, have been suggested as alternative solutions, which can be deployed dynamically and on physiological time scales. A number of *in vitro* studies on archaeal histones (Sandman *et al*. 1990), Sul7 (Baumann *et al*. 1994; Guagliardi *et al*. 1997), HTa (Stein and Searcy 1978; Searcy 1986), and MC1 (Chartier *et al*. 1988) have shown that NAP binding can significantly increase melting temperature, reduce the risk of DNA denaturation, and/or promote strand re-annealing. The results reported here unify these findings to establish temperature as a universal quantitative driver of investment in NAPs across the archaea.

NAPs likely play a role in reducing the risk of accidental as well as programmed opening events, which occur in the context of transcription, replication, and repair. A particular risk in this regard emanates from promoters, which are AT-rich and poised to open to enable transcription. Thermophiles appear to have reduced this liability in part through a simple strategy: losing promoters. The number of genes per transcription unit, co-expressed from a single upstream promoter, increases with temperature (Fig S12). In addition, the proportion of the genome dedicated to intergenic regions decreases with temperature across archaea (Sabath *et al*. 2013). Pinning the promoter on either side with DNA-binding proteins – as observed for histones (Nalabothula *et al*. 2013) but also HTa in *T. acidophilum* (Hocher *et al*. 2019) – might have evolved in parallel to prevent uncoordinated promoter melting and runaway extension of the resulting denaturation bubbles.

Protection from denaturation as the principal function of NAPs is consistent with the high fluidity of NAPs across archaeal evolution, epitomized by the Diaforarchaea. Proteins of several different folds will probably do a serviceable job of raising DNA melting temperature and curbing denaturation if they found themselves transplanted into a different genomic context. This model of NAP evolution is further consistent with limited quantitative covariation between the abundance of NAPs and other chromatin factors across evolution and with NAPs being encoded as single-gene operons. Both observations suggest a scarcity of intimate dependencies.

At the same time, we must highlight that our results do not exclude specific adaptive roles that may drive NAP diversity across the archaea. One such adaptive role might be in the prevention, detection, and repair of DNA damage (Grogan 1998). Mutagenic challenges differ considerably across environments and might favour some NAPs over others. MC1, for example, protects against radiation damage (Isabelle *et al*. 1993), a frequent insult for halophiles that live in shallow aquatic environments. Conversely, Cren7 binds to T:G mismatches produced by cytosine deamination events (Tian *et al*. 2016), which become more common at higher temperature. Thus, we suggest that while denaturation concerns shape total NAP abundance, NAP diversity is likely a product of both exchangeability and species-specific requirements for nucleoid function and maintenance.

New chromatin components can be acquired from other archaea or bacteria, as illustrated by the eventful natural history of the Diaforarchaea where HTa, Alba, MC1 and the newly identified Methanomassiliicoccales protein can trace their origin to horizontal transfers. In addition, new components can arise through innovation/repurposing of proteins already present in the proteome, as transcription factors, like TrmBL2 in *T. kodakarensis*, become global chromatin constituents. Conversely, proteins can lose their global architectural roles. In the most extreme case, once abundant NAPs can be lost entirely – as observed in the Methanomethylophilaceae. They can also undergo significant reductions in abundance. This is what appears to have happened to histones in halophiles and other lineages, consistent with their non-essential status in *H. salinarum* (Dulmage *et al*. 2015) and *Methanosarcina mazei* (Weidenbach *et al*. 2008). The low abundance of HstA in *H. volcanii*, at both the transcript (Rojec *et al*. 2019) and protein level (Fig 3D), is hard to reconcile with its purported role as major architectural factors (Ammar *et al*. 2011). We therefore suggest that prior findings of widespread protection from micrococcal nuclease digestion in this species might, in fact, be caused not by histones but by an as yet uncharacterized protein or set of proteins. Our *de novo* prediction pipeline suggests HVO_1577, a protein that contains an HrcA DNA-binding domain, as a candidate. Quantitative chromatin mass spectrometry and/or histone/HVO_1577 deletions in *H. volcanii* will help to clarify this issue.

Our data show that, when moving from a thermophilic to a mesophilic niche, different lineages of archaea have reduced their investment in NAPs – shedding a cost that can now be spared. Does this imply that all archaeal mesophiles exhibit reduced investments in chromatin? We suspect not. For example, histones are very highly expressed, at least at the transcript level, in some mesophilic members of the Methanobacteriales, notably *Methanobrevibacter smithii*, which grows at 37°C (its three histones are ranked 1^st^, 8^th^, and 282^nd^ most highly expressed) (Rojec *et al*. 2019). Whether this also holds true at the protein level remains to be established, but we suggest that high levels of chromatinization might be obligatory for thermophiles but facultative for mesophiles.

Finally, eukaryotes are (in the vast majority) mesophiles, yet their DNA is ubiquitously wrapped in nucleosomes and histones have become a linchpin whose removal leads to the collapse of controlled gene expression (Hennig and Fischer 2014). The acquisition of histone tails and their subsequent use for signalling might have been one of the factors driving entrenchment, generating a thick top layer of cellular machinery that acts on, modifies, and remodels nucleosomes to orchestrate gene expression, DNA repair and replication. Over time, the evolution of cryptic promoters, rendered inaccessible by nucleosomes but activated following their removal, might also have contributed to the retention of global chromatinization (Hennig and Fischer 2014). Irrespective of the factors that first rendered eukaryotic histones indispensable, we speculate that high levels of chromatinization in eukaryotes might represent an evolutionary hangover from their thermophilic ancestry, and that eukaryotes – unlike many of their archaeal cousins – evolved a dependency on global chromatinization that they were unable to break when moving into a more temperate niche.

## METHODS

### Culture of Methanomassiliicoccales

*M. luminyensis* was obtained from DSMZ (DSM 25720) and *M. alvus* (isolate Mx-05) had been isolated previously by one of us (JFB). Both strains were grown in strictly anaerobic conditions (2 atm. of H_2_/CO_2_ – 20/80) using 10 mL of growth medium in 50 mL glass bottles according to DSMZ recommendations for *M. luminyensis* with one exception: ruminal fluid (200 µL) was added for *M. alvus*. Cultures were performed without shaking at 37°C using 60 mM of methanol as electron acceptor for methanogenesis.

### Protein pellet preparation and mass spectrometry

50 mL aliquots of 10-day (3-day) cultures of *M. luminyensis* (*M. alvus*) were pelleted with care to preserve anoxic conditions as much as possible and stored at -80°C. Protein pellets were prepared following PreOmics iST kit guidelines. 10mg of frozen pellet were resuspended in lysis buffer and volumes adjusted after the heating step to load 100µg as measured by nanodrop absorbance at 205nm. A DNA sonication step was included (Diagenode Bioruptor) followed by digestion for 1.5 hrs. The whole procedure was carried out without interruption and pellets stored at -80°C in MS-LOAD buffer before being processed by mass spectrometry (two biological replicates with technical replicates for each).

### Genomes database

All genomes and proteomes were obtained from NCBI assembly (https://www.ncbi.nlm.nih.gov/assembly) accessed on 2021-05-21. Proteomes that were not available from NCBI were predicted using Prodigal v2.6.3 using default parameters.

### Species tree and taxonomy

The archaeal species tree and taxonomic groups were obtained from GTDB (https://gtdb.ecogenomic.org) accessed on 2020-09-23, with some species names updated to reflect current use in the literature (Table S1).

### Processing of public proteomics data

We only included proteomes that were derived a) from whole cell extract, b) without size selection and c) comprised more than 500 identified proteins. Data for *H. volcanii* was obtained from Jevtić *et al*. (2019) and (for Fig S7) Knüppel *et al*. (2021), *T. kodakarensis* from Sas-Chen *et al*. (2020), *N. magadii* from Cerletti *et al*. (2018), and *N. maritimus* from Qin *et al*. (2018). For each dataset, measurements that did not correspond to the Genbank complete genome of the strain/species were discarded. Correspondence between Uniprot and Genbank ID was established using the Uniprot Retrieve/ID mapping tool. For each dataset, normalized intensity (in %) was computed as the ratio of each protein intensity over the total intensity for the species of interest.

### Protein annotations

HMM models were downloaded from PFAM (PFAM-A, accessed on 2020-01-20) and TIGR (TIGRFAMs 15.0, accessed on 2020-05-15) and sequences were searched using hmmsearch (version 3.1b2). All searches were carried out using the gathering thresholds provided for each models (option -ga) to ensure reproducibility. No further threshold was applied unless mentioned otherwise. Results were robust to application of an alternative, stricter threshold of 1e-3. As no HMM model existed for C1, we searched for sequences homologous to *Thermoproteus tenax* Cc1 (Uniprot ID : G4RKF6) using jackhmmer (version 3.1b2), applying an e-value threshold of 1e-5. A list of DNA-binding protein PFAM domains was obtained from (Malhotra and Sowdhamini 2015) and domains found in transcription factors were manually annotated from this list. Detailed tables and full sequences of all NAPs and candidate NAPs discussed in this study are available as supplemental material (Table S1).

### Gene ontology

Gene ontologies were obtained from pfam2go tables, available at http://current.geneontology.org/ontology/external2go/pfam2go. When computing correlations between environmental variables and gene ontology, each protein was only counted once per ontology.

### Habitat and phenotypes

Phenotypic data was obtained from Madin *et al*. (2020) and habitat data from ProGenome Mende *et al*. (2020). When multiple strains per species or multiple sources per strain were available, optimal growth temperature was averaged, with the exception of *H. volcanii* for which the average was seemingly too low (38°C) and was thus set at 45°C, based on the description of the original isolate. Growth rates for *M. luminyensis* and *M. alvus* (absent from the database) were added manually.

### Operon prediction

Operons were predicted using Operon Mapper (https://biocomputo.ibt.unam.mx/operon_mapper/) with default settings.

### DNA binding prediction

DNA binding was predicted using DNAbinder (https://webs.iiitd.edu.in/raghava/dnabinder/) using the SVM model trained on a realistic dataset. Proteins whose score was higher than 0 were considered a possible DNA binding proteins.

### Protein alignments and phylogenetic trees

Proteins sequences were aligned using MAFFT (option -linsi). With the exception of the species tree (see above), all trees were built using RAXML-NG, model LG+R6. Best maximum likelihood midpoint rooted trees are shown along with the results of 100 non-parametric bootstraps. Trees were visualized using iTol (https://itol.embl.de/).

### Phylogenetic linear regression

To control for phylogenetic non-independence, phylogenetic linear regression were carried using the R package phylolm, Model “BM” with 10000 bootstraps or ‘OUrandomroot’. Variables were log transformed before regression.

### Scripts

All scripts are available at https://github.com/hocherantoine/NAPsQuant

## ACKNOWLEDGEMENTS

We thank Holger Kramer and the LMS Proteomics Facility for generating mass spectrometry data and the Molecular Systems Group for discussions. This work was funded by Medical Research Council core funding (TW), an EMBO Short-Term Fellowship 8472 (AH) and the French National Agency for Research (SG; Grant ArchEvol ANR-16-CE02-0005-01).

## SUPPLEMENTARY DATA

**Table S1**. Known and candidate NAPs across 1419 archaeal genomes.

**Table S2**. Correlations between NAP abundance and ecological covariates.

**Figure S1.**
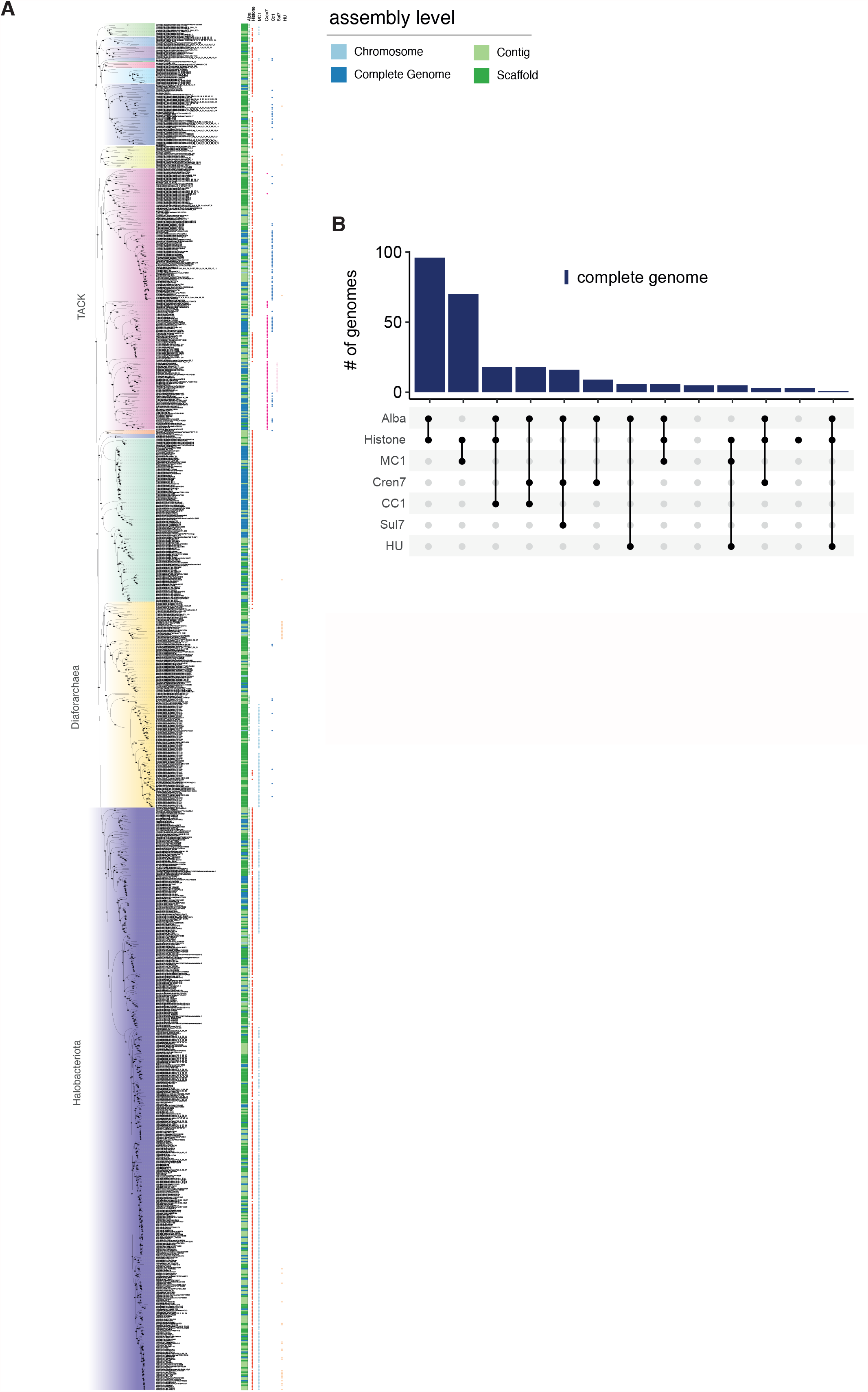
Distribution of nucleoid-associated proteins (NAPs) across the archaea. (**A**) Name and assembly level are provided for each species in phylogenetic context. The species-level phylogeny is based on GTDB (see Methods). (**B**) Co-occurrence of NAPs. This panel is equivalent to Fig 1B but only considers co-occurrence in complete genomes. Information on assembly level is based on metadata from NCBI.

**Figure S2.**
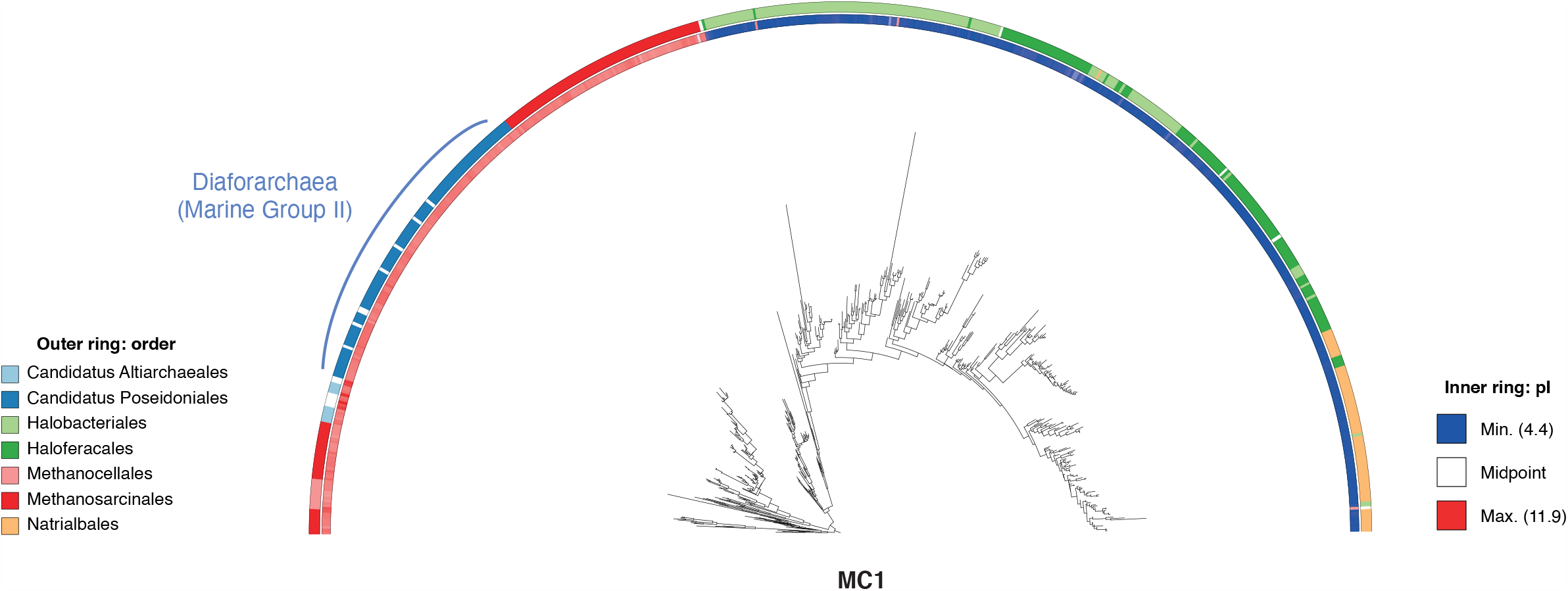
Maximum likelihood phylogeny of archaeal MC1 homologs. MC1 sequences found in the Diaforarchaea are restricted to marine group II archaea and branch as a monophyletic group within the Methanosarcinales consistent with a single horizontal transfer event. Note also that sequences from haloarchaea have dramatically lower isoelectric points, making them unlikely donors.

**Figure S3.**
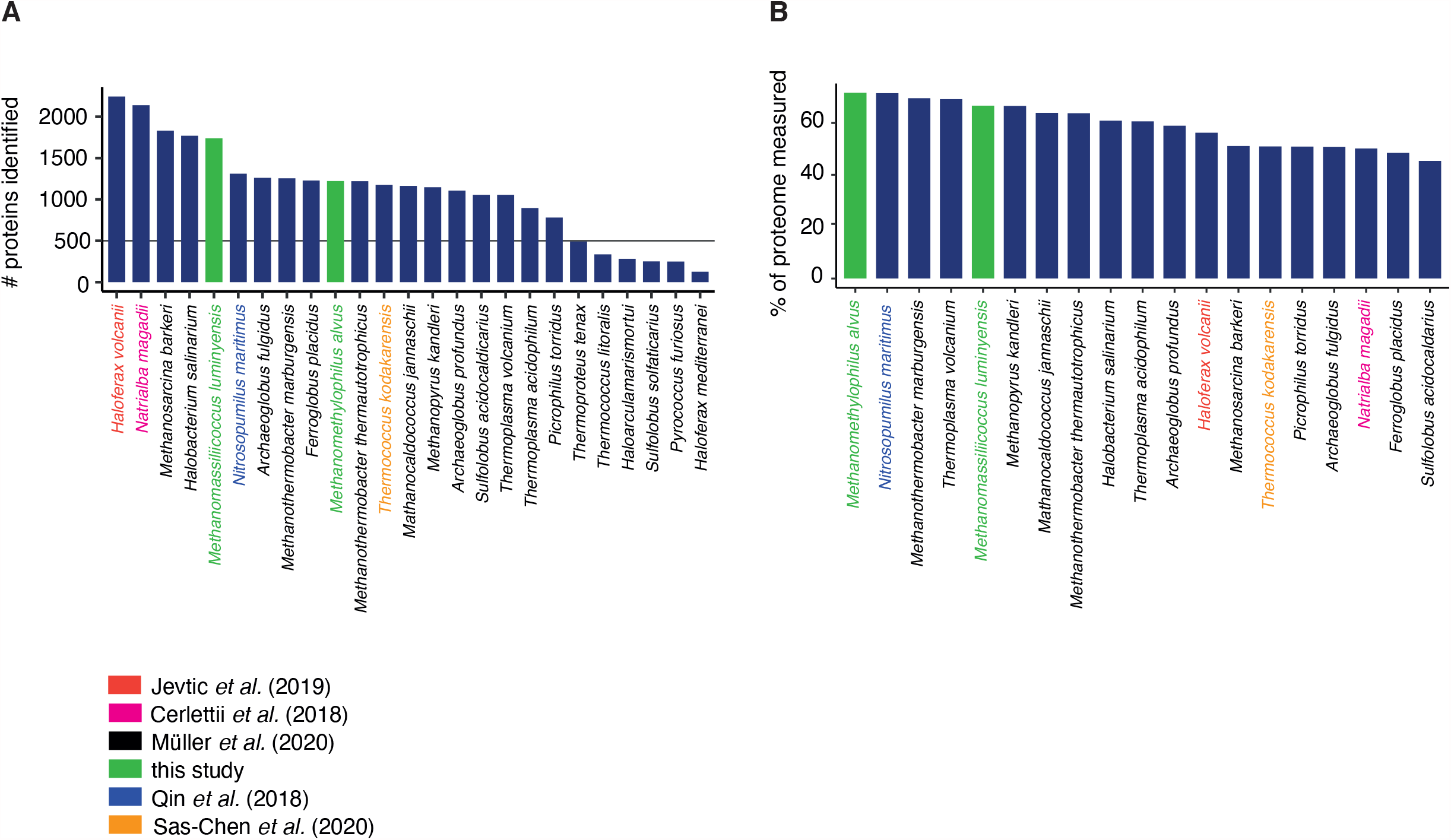
Absolute (**A**) and relative (**B**) coverage of the predicted proteome in 19 archaea.

**Figure S4.**
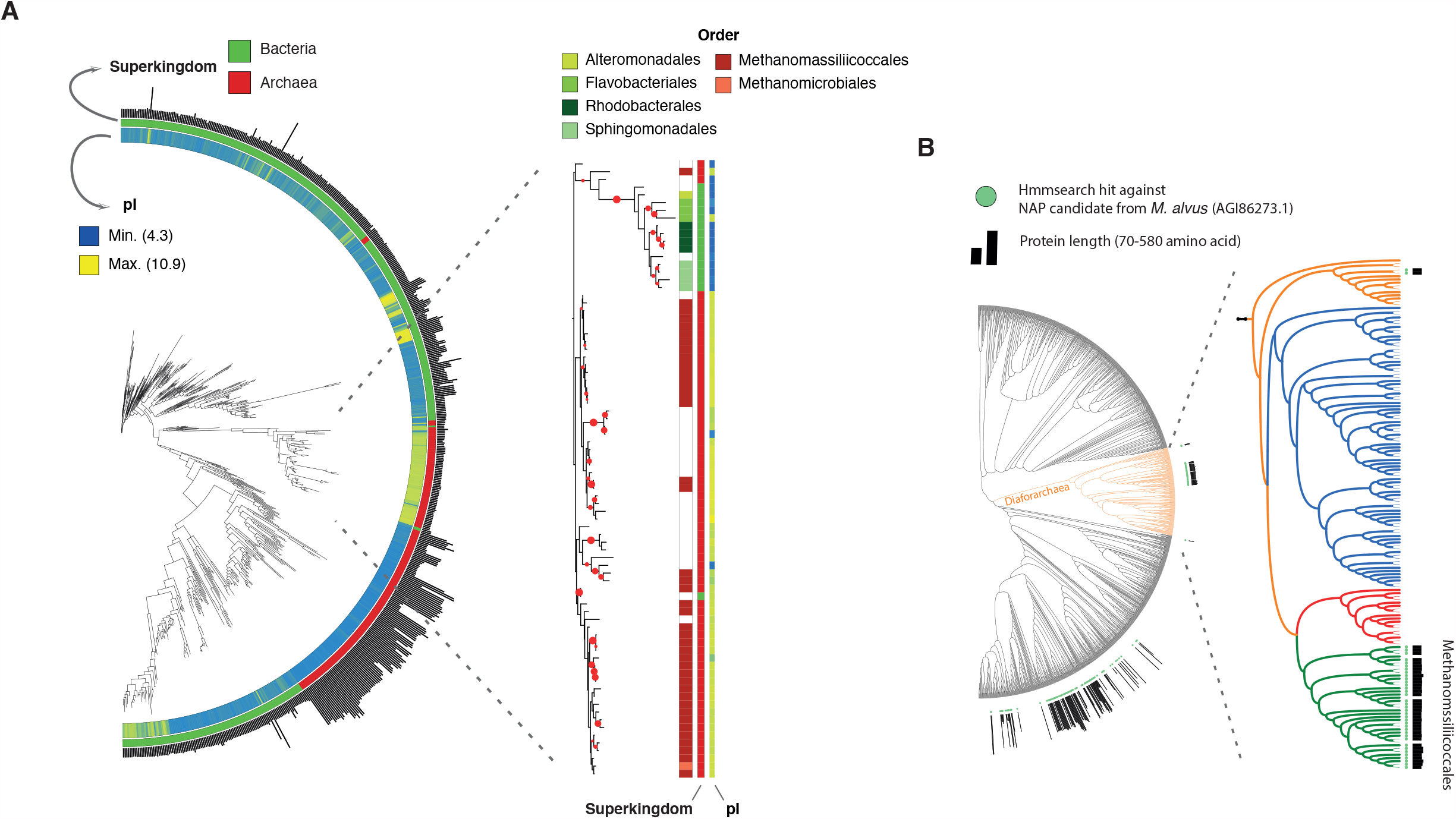
The novel candidate NAP WP_019177984.1/AGI86273.1 in phylogenetic context. (**A**) Maximum likelihood phylogeny of bacterial and archaeal WP_019177984.1/AGI86273.1 homologs. Homologs were obtained by running jackhmm using AGI86273.1 as a seed (e-value 1e-5). The subtree indicated by dashed lines was re-aligned and a tree computed using RAxML-NG. Bootstrap (n=100) values superior to 50 are shown. (**B**) Distribution of jackhmm hits using AGI86273.1 as a seed across archaea, displayed on a species tree obtained from GTDB.

**Figure S5.**
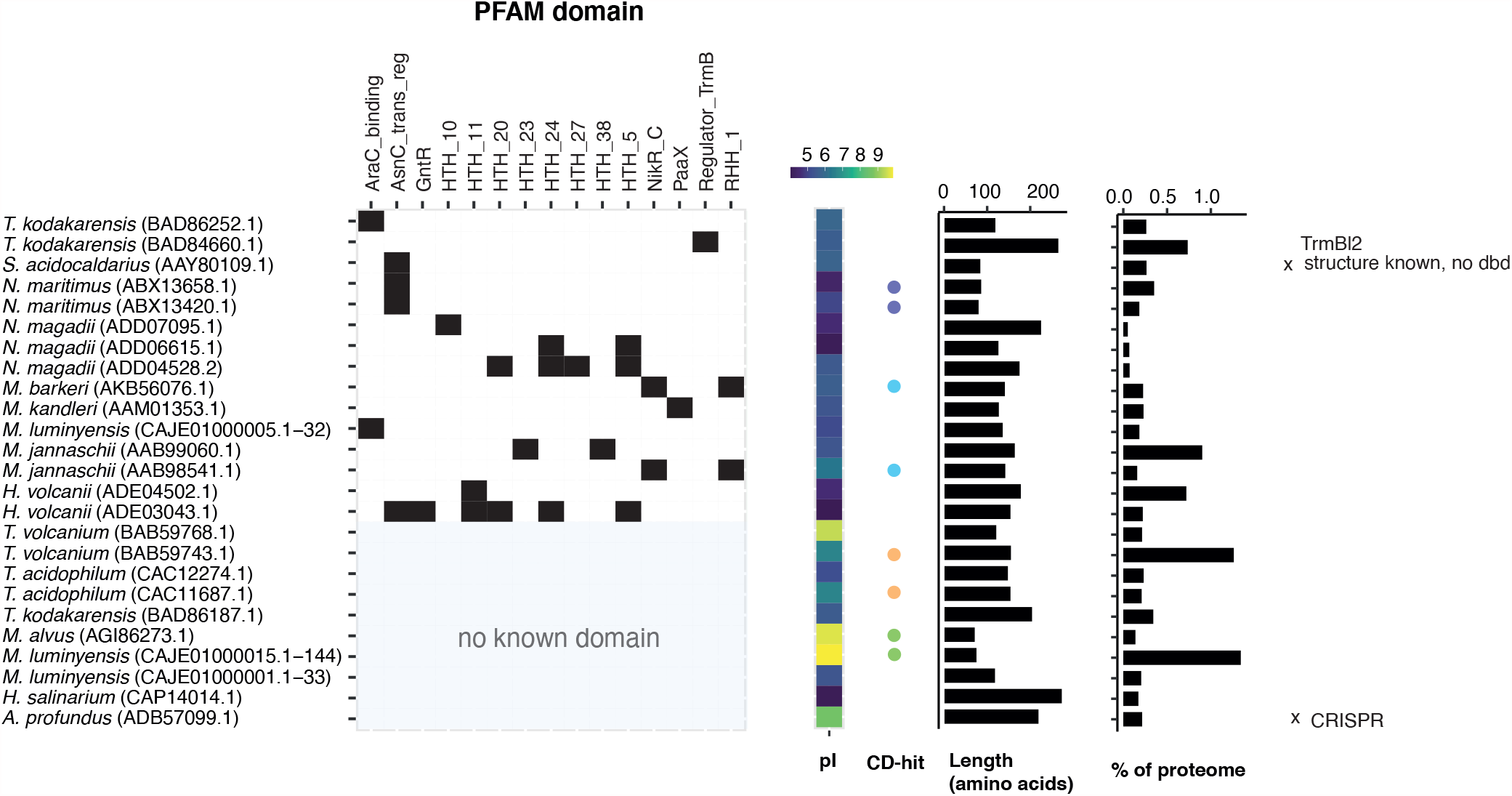
Overview of candidate NAPs, their domain content, size, isoelectric point (pI) and fractional abundance in the proteome of the species in which they were identified. Likely orthologs amongst this set were identified using CD-hit (see Methods) and bear the same colour. Two likely false positives are marked with an “x”. The pipeline used to predict novel candidate NAPs is described in the main text and Methods. Also see Table S1 for further details.

**Figure S6.**
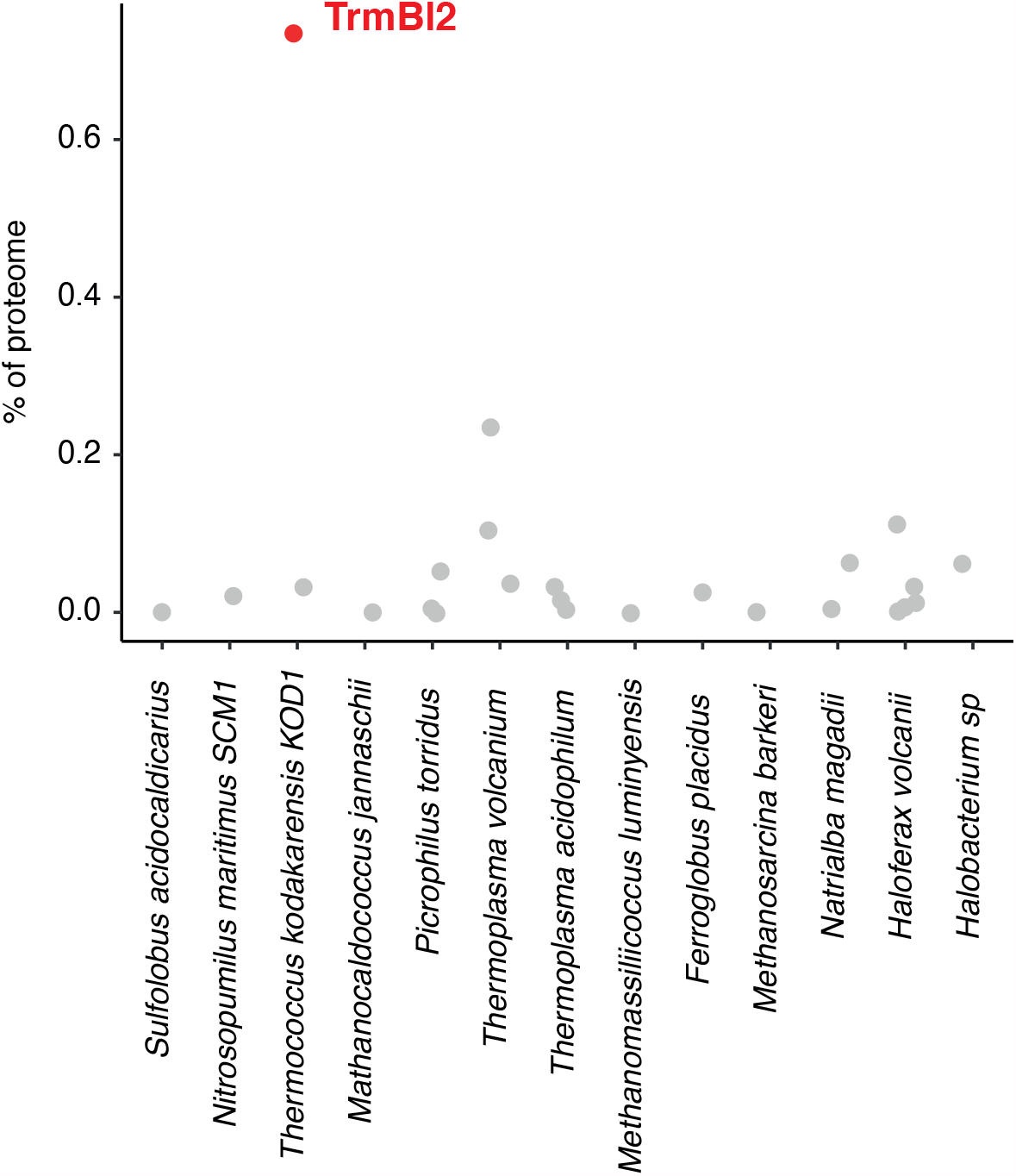
Relative abundance of TrmBL orthologs in different proteomes. *T. kodakarensis* TrmBL2 stands out as having a much higher relative abundance than its homologs in other proteomes.

**Figure S7.**
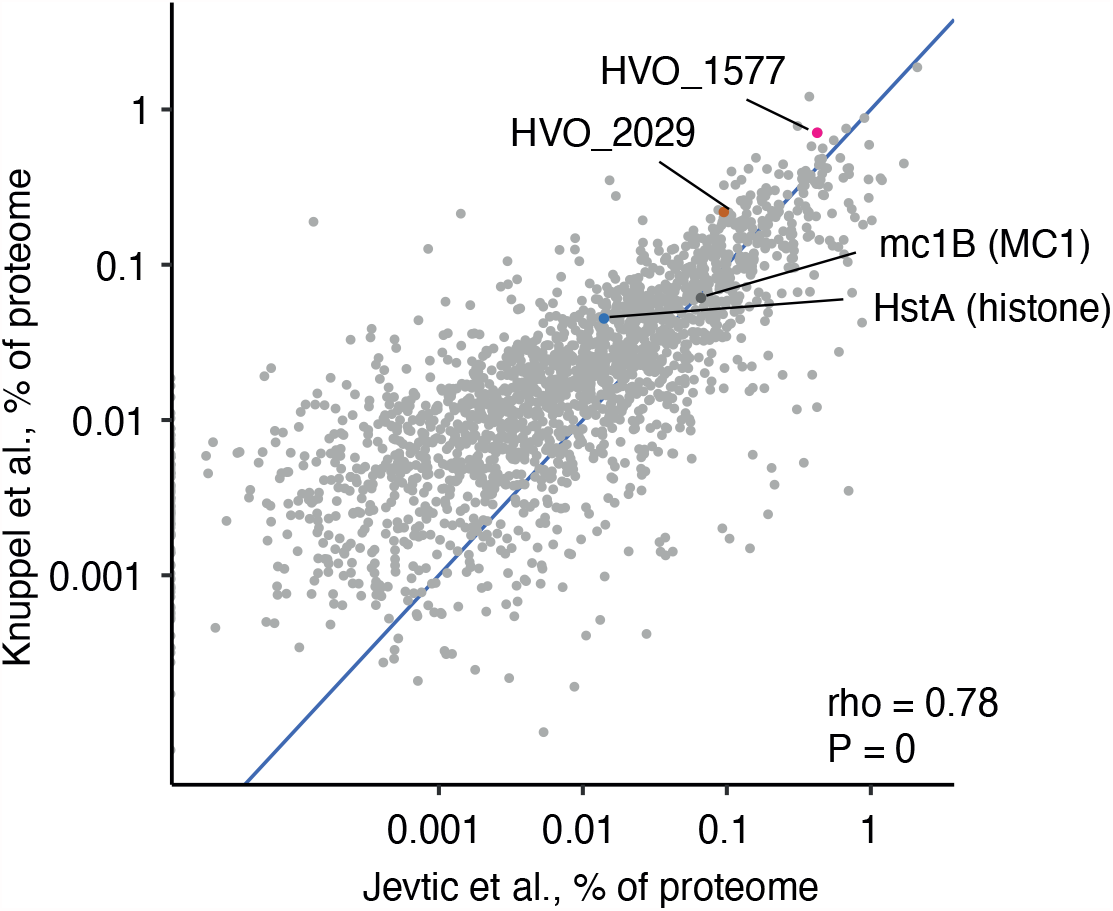
Relative abundance of known and candidate NAPs in *Haloferax volcanii*, as determined in two other quantitative proteomics (Jevtić *et al*. 2019; Knüppel *et al*. 2021). Note that the single *H. volcanii* histone (HstA) and MC1 are >10-fold less abundant than the novel NAP candidate HVO_1577.

**Figure S8.**
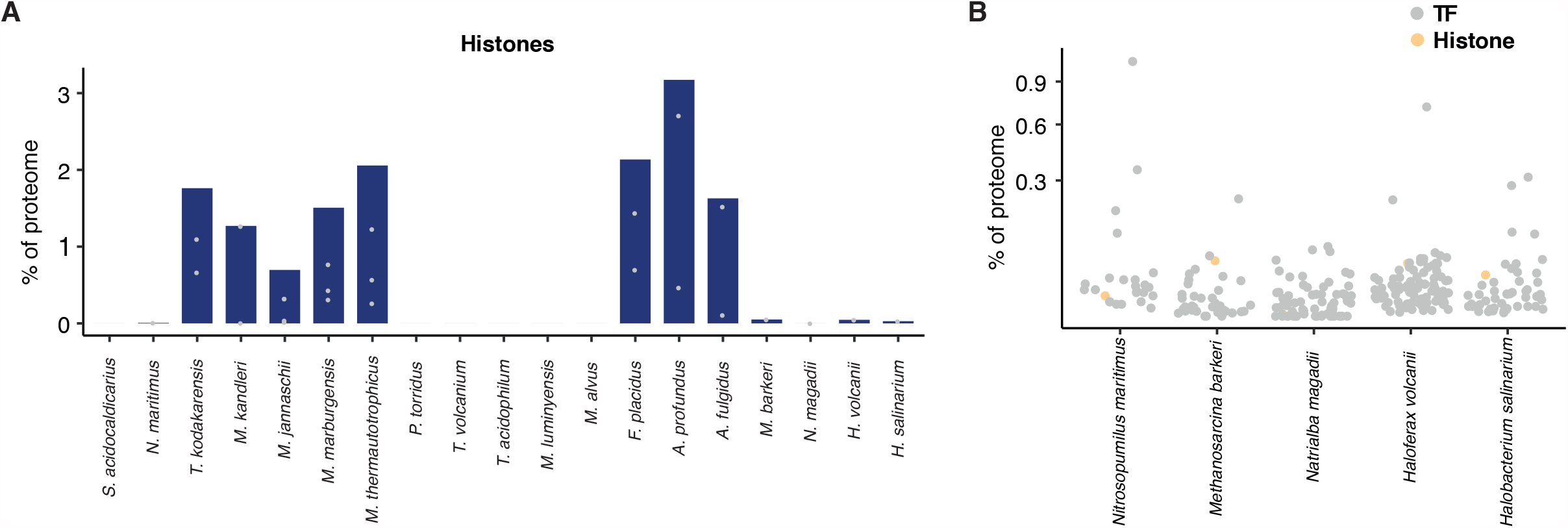
Quantitative variation of histone abundance across 19 archaea. (**A**) Bar heights represent the summed abundance of all detected histone paralogs in a given species, while the grey dots mark the abundance of individual paralogs. (**B**) The relative abundance of histone proteins in halophilic archaea compared to the abundance of transcription factors (TF) measured in the same experiment.

**Figure S9.**
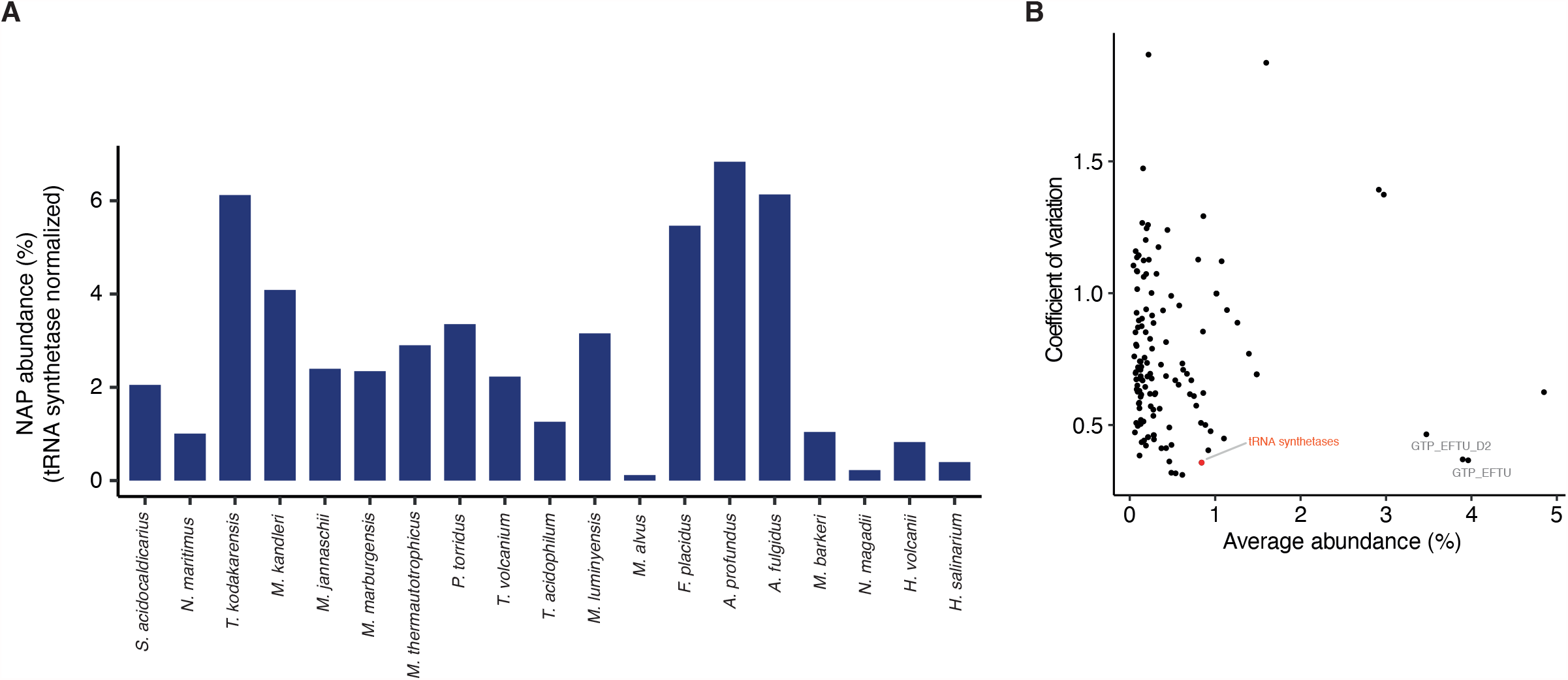
(**A**) Relative NAP abundance normalized to the abundance of tRNA synthetases in the same proteome. (**B**) Compared to other Pfam domains, proteins classified as tRNA synthetases are both highly abundant and different organisms dedicate a similar fraction of their protein budget to their production (as evident from a low coefficient of variation across species), making them a suitable choice for normalization. The idea of normalization here is to consider the investment in NAPs relative to some baseline investment in basic cellular processes (such as charging amino acids). The results indicate that large quantitative variability in NAP abundance is also evident when Sjudged against this baseline rather than as a fractional allocation across the entire proteome.

**Figure S10.**
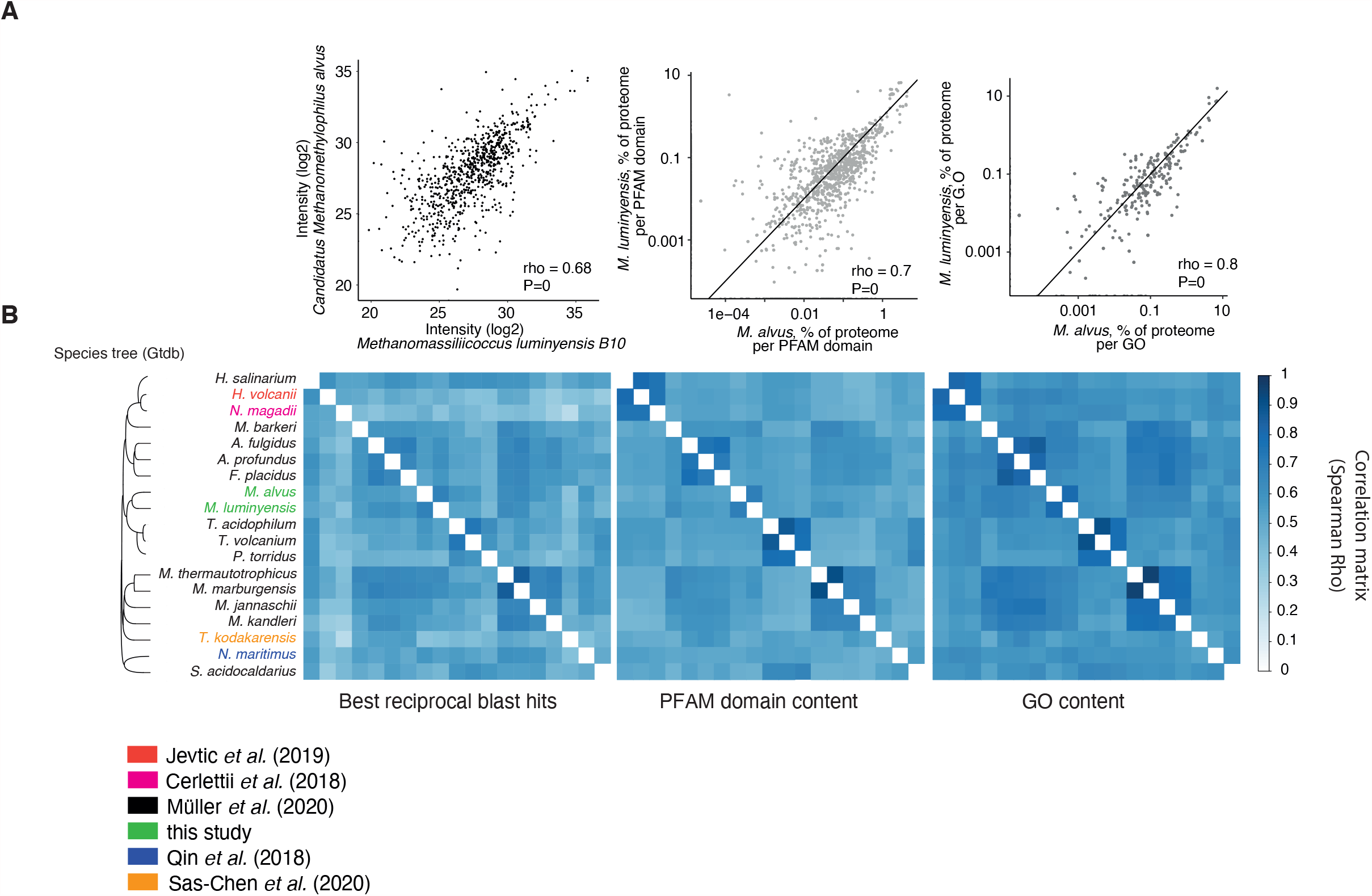
Comparing relative protein abundances across proteomes. (**A**) Correlation of relative protein abundances in *Methanomassiliicoccus luminyensis* versus *Methanomethylophilus alvus* when comparing reciprocal best BLAST hits (left panel) or proteins aggregated by Pfam domain content (middle panel) or by gene ontology class (right panel). (**B**) Visualization of pairwise correlation coefficients for the same comparisons across all 19 proteomes. The species tree is taken from GTDB, with *P. torridus* and *H. salinarum* added manually.

**Figure S11.**
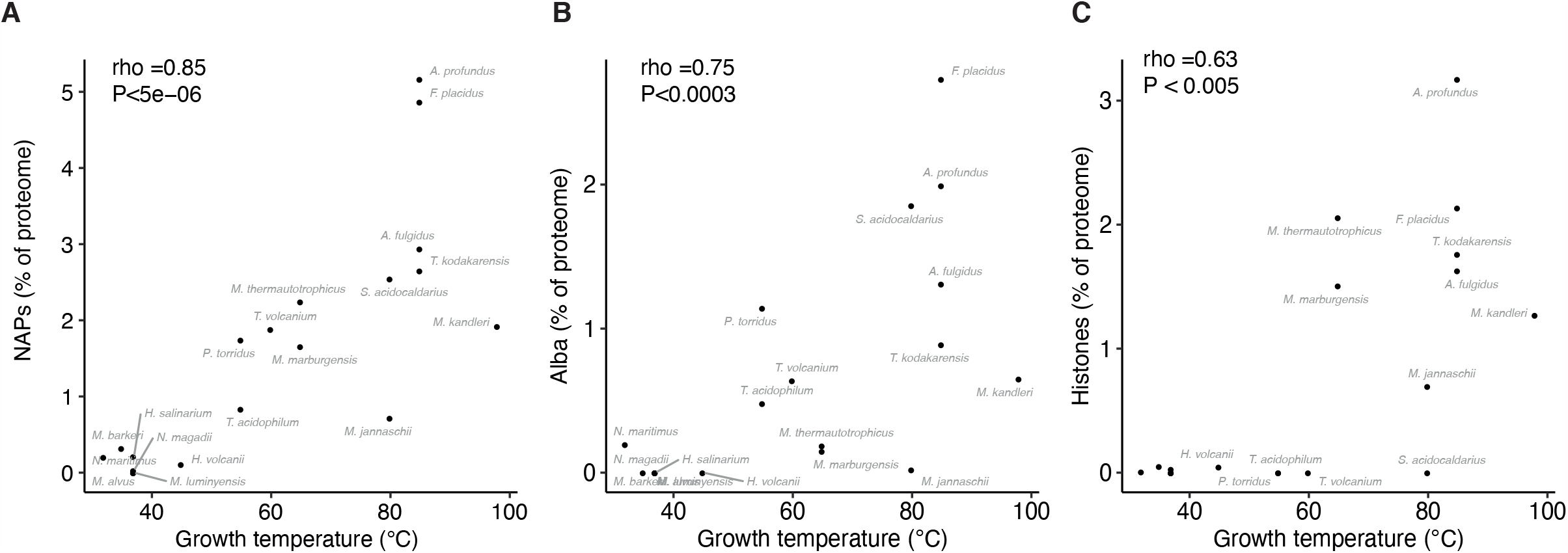
Relative abundance of individual NAPs as a function of optimal growth temperature. Strong correlations are evident not only for (**A**) NAPs considered as an aggregate class but also for (**B**) Alba and (**C**) histones considered individually. Note that Alba and histones are the only NAPs that are sufficiently widespread across our sample of archaea to allow meaningful correlations to be computed.

**Figure S12.**
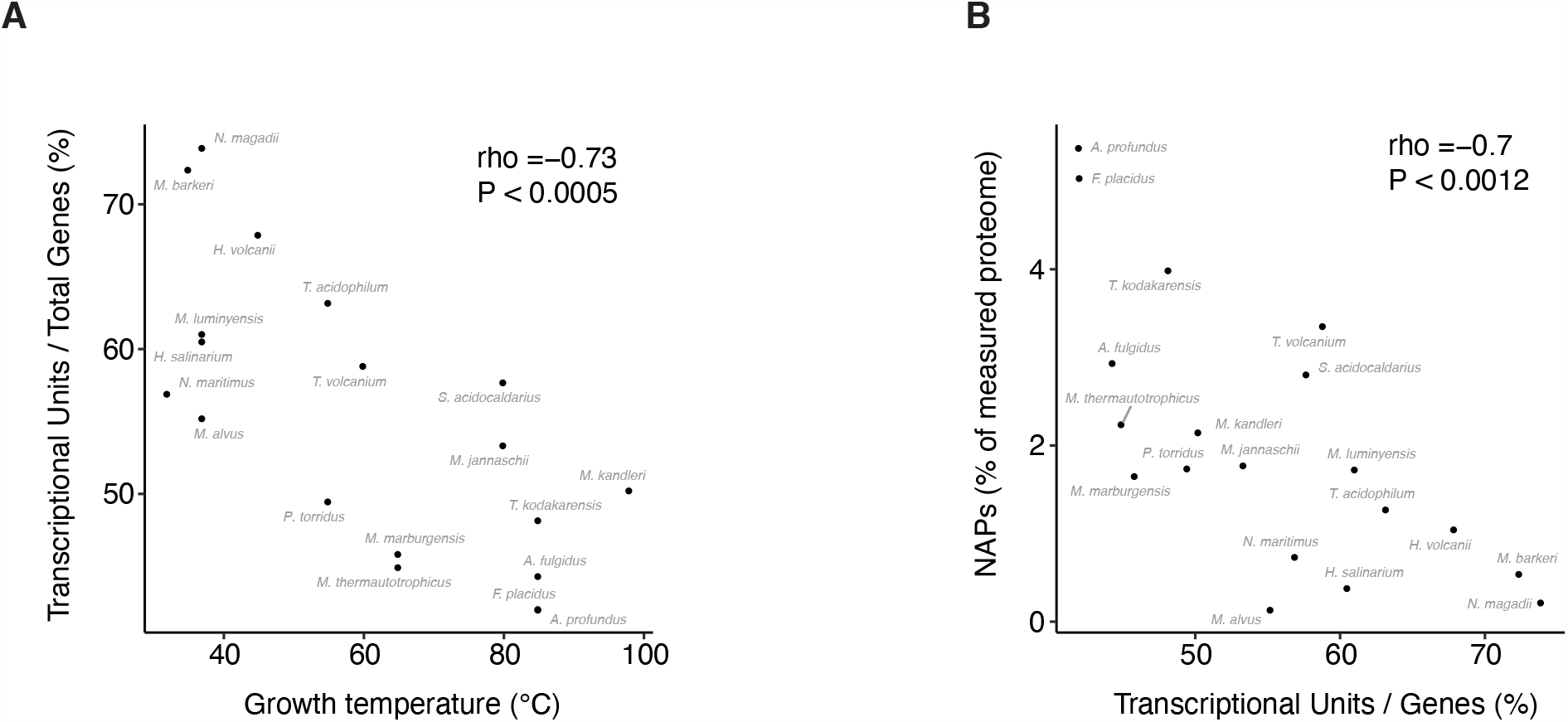
The relationship between growth temperature and genome compactness. (**A**) The genomes of organisms growing at higher temperatures tend to be organized in such a manner that transcript units harbour more genes on average. There are therefore fewer promoters per gene in archaea that grow at higher temperature. (**B**) As predicted from the respective relationships with optimal growth temperature, genomes where genes are contained in more independent transcriptional units have a lower investment in NAPs.

